# Modeling subcortical white matter stimulation

**DOI:** 10.1101/2019.12.12.872390

**Authors:** Melissa Dali, Jennifer S. Goldman, Olivier Pantz, Alain Destexhe, Emmanuel Mandonnet

## Abstract

**Objective:** Intracranial electrical stimulation of subcortical axonal tracts is particularly useful during brain surgery, where mapping helps identify and excise dysfunctional tissue while avoiding damage to functional structures. Stimulation parameters are generally set empirically and consequences for the spatial recruitment of axons within subcortical tracts are not well identified.

**Approach:** Computational modeling is employed to study the effects of stimulation parameters on the recruitment of axons: monophasic versus biphasic stimuli induced with monopolar versus bipolar electrodes, oriented orthogonal or parallel to the tract, for isotropic and anisotropic tracts.

**Main results:** The area and depth of axonal activation strongly depend on tissue conductivity and electrode parameters. The largest activation area results from biphasic stimulation with bipolar electrodes oriented orthogonal to axonal fasciculi, for anisotropic and especially isotropic tracts. For anisotropic tracts, the maximal activation depth is similar regardless of whether a monopolar or bipolar electrode is employed. For isotropic tracts, bipolar parallel and monopolar stimulation activate axons deeper than orthogonal bipolar stimulation. Attention is warranted during monophasic stimulation: a blockade of action potentials immediately under cathodes and a propagation of action potentials under anodes are found.

**Significance:** Considering the spatial patterns of blockade and activation present during monophasic stimulation with both monopolar and bipolar electrodes, biphasic stimulation is recommended to explore subcortical axon responses during intraoperative mapping. Finally, the precise effect of electrical stimulation depends on conductivity profiles of tracts, and as such, should be explicitly considered for each individual subject and tract undergoing intracranial mapping.

## Introduction

Intra-operative functional brain mapping is performed during tumor excision, while patients are awake, to interrogate the organisation and operation of tissues. As such, functional mapping aids in the identification of non-operational tissues important to excise, and the prevention of neuropsychological complications induced by the excision of operational tissue and tracts (fasciculi of axons). Despite the demonstrated utility of intra-operative functional mapping, there is no standard approach to choose the electrode parameters that are mostly set empirically [31]. Parameters of electrical stimulation include pulse shape, pulse duration (*PW*), stimulus amplitude (*I*), (Fig. 1), stimulation frequency, as well as the polarity and orientation of electrodes. To avoid tissue damage and ensure safe stimulation, the net charge injection (ie. the stimulus amplitude *I* multiplied by the stimulus duration *PW*) must be equal to zero [19]. This condition can be reach using biphasic pulse (Fig. 1 C) instead of monophasic pulse (Fig. 1 A and Fig. 1 B). Although bipolar biphasic stimulation is often used during subcortical or cortical electrical stimulation [31], some clinical studies have also explored cortical and subcortical responses using bipolar monophasic [13, 30], monopolar monophasic [10], and monopolar biphasic stimulation [10]. The effects of stimulation parameters are difficult to assess experimentally, because closed-loop electrodes are not routinely used for functional mapping and other fine assessments of the spatial-temporal effects of stimulation parameters would require implantation of further electrodes used only for research, and thus difficult to pass ethical review. Another issue is the variability of electrical conductivity in the white matter [26]. Previous studies have shown that the tissue anisotropy surrounding the electrode can alter the shape of the electric field and the subsequent neural response to stimulation [9, 28]. To study the effects of stimulation parameters despite empirical limitations, computational modeling of neural responses to stimulus-induced electric fields are commonly used, especially in deep brain stimulation [2, 18, 20, 27, 33] but few have studied the effects of subcortical stimulation [8, 16, 17]. Gomez-Tames et al. [8] have recently assessed the influence of electrode diameter, and inter-electrode distance, as well as bipolar versus monopolar electrode using a head model with purely isotropic white matter. They show that for a fixed current amplitude, monophasic monopolar stimulation had a broader activation region that monophasic bipolar stimulation. The orientation of the bipolar electrode according the axonal tract was not taken into account. The studies of Mandonnet and Pantz [16, 17] have shown that, using a biphasic pulse, a bipolar electrode oriented orthogonal to the axonal tract allowed broader activation of the tract than an electrode oriented parallel. Although subcortical modeling studies have contributed significantly to understanding the effects of stimulation parameters, the pulse shape variability as well as the influence of the axonal tract anisotropy with respect to the polarity and orientation of electrodes have not been yet investigated.

**Figure 1.**
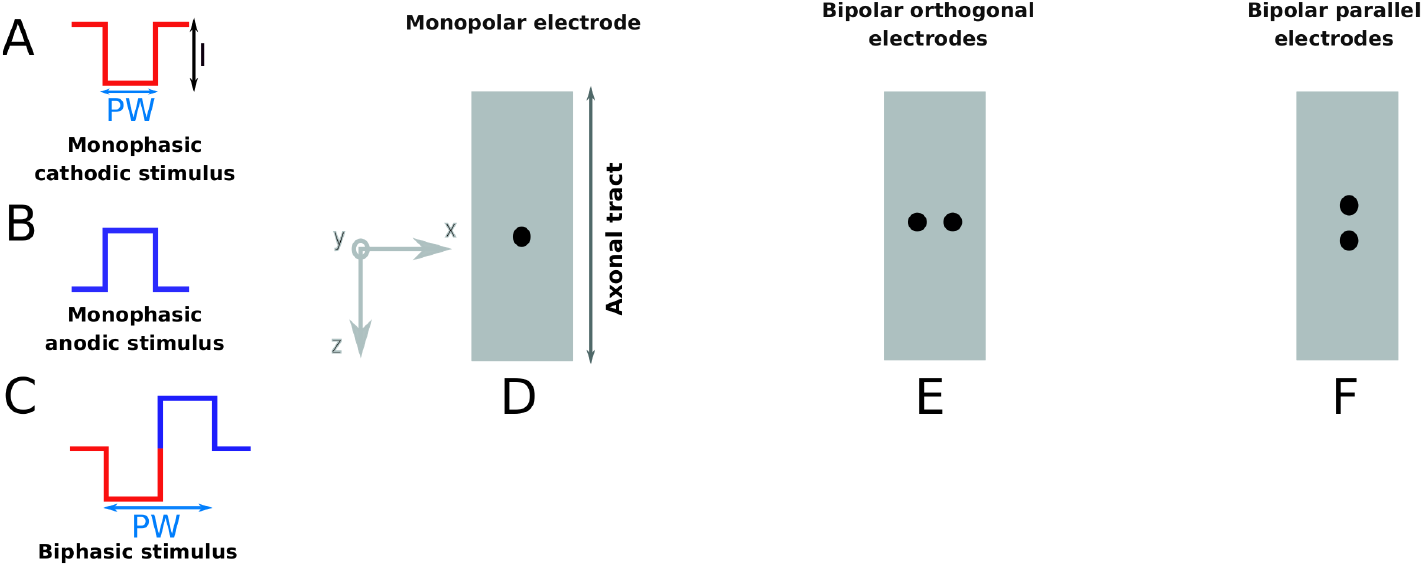
(A-C): Stimulus parameters. (A): monophasic cathodic (depolarizing) stimulus, (B): monophasic anodic (hyperpolarizing) stimulus, (C): biphasic stimulus, (D-F): electrode configurations (represented by black dots), longitudinal view of an axonal tact. (D): monopolar electrode, (E): bipolar orthogonal electrodes, (F): bipolar parallel electrodes

The present study aims to deepen the knowledge of the effects of stimulation parameters used to evoke subcortical responses. For this purpose, a model was developed to compute the response of a myelinated axons of a tract (either isotropic or anisotropic) to the potential field generated by various electrode parameters: monophasic versus biphasic pulses, through monopolar versus bipolar electrodes, oriented parallel versus orthogonal to tracts.

## Material and methods

### Two-part model

A volume conductor model for an axonal tract and bipolar (or monopolar) electrodes was coupled with a model of mammalian myelinated nerve fiber to study axonal recruitmentment resulting from stimuli. This method builds on models developed to successfully describe the effects of peripheral nerve stimulation [4, 5], adapted for white matter stimulation of tracts in the central nervous system.

### Volume conductor model

We considered a bundle of axons enclosed in a half-cylindrical model (10 *mm* diameter) embedded in a sphere representing the surrounding white matter. The 3D FEM model was implemented on COMSOL Multiphysics (COMSOL Inc, Burlington, MA) software. Three models of electrodes were considered (Fig. 1, D-F): monopolar with distant reference (10 cm from the stimulating electrode), bipolar oriented parallel to the axonal tract, and bipolar oriented orthogonally to the axonal tract. The inter-electrode distance was 7 *mm* and the electrode diameter, 1 *mm* [16, 17]. First, the fasciculus and the surrounding tissue were considered entirely isotropic with conductivity *σ*_*iso*_ = 0.14 *S*/*m* [6]. Then, the axonal tracts were considered anisotropic. We used the volume constraint method [11, 32] to compute the values of the conductivity tensor.

This method retains the volume between the anisotropic and the corresponding isotropic tensor:

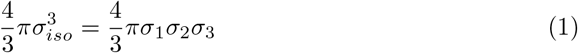

*σ*_*iso*_ is the isotropic conductivity of the white matter, *σ*_1_, *σ*_2_ and *σ*_3_ are the eigenvalues of the conductivity tensor. Eq. Eq. 1 was parameterized in terms of two ratios [11]:

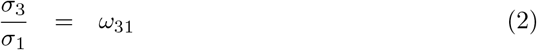

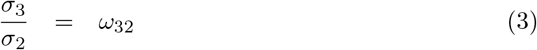

Using Eq. 1, 2, and 3; *σ*_1_, *σ*_2_ and *σ*_3_ were expressed as follow:

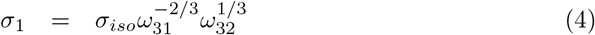

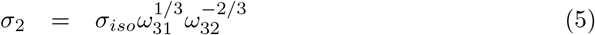

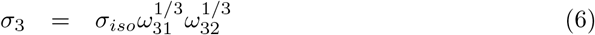

The *σ*_3_ eigenvector was set as 9 times larger than the values of the perpendicular eigenvectors: *ω*_31_ = *ω*_32_ = 9 [21]. The isotropic white matter conductivity was also applied to the surrounding tissue.

The potential fields through the volume conductor model resulting from electrode stimulation, were computed following the same method described in previously [5]. Briefly, the Poisson equation was solved using COMSOL, assuming quasi-static conditions [1] and appropriate Neumann Dirichlet boundary conditions [22].

### Axon model

The field simulation was then coupled to non-linear cable models of myelinated axons. The diameter size was fixed to 10 *μm*, consistent with the diameter of myelinated axons (considering both axon and myelin) found in the central nervous system [14]. Spacing in the radial direction (X,Y) was set to 100 *μm*. Spacing on longitudinal direction (Z), the internodal spacing, was set to 1000 *μm* [12]. Models of mammalian myelinated fibers were implemented in Matlab (The MathWorks, Natick, Massachusetts).

Chiu-Ritchie-Rogart-Stagg-Sweeney (CRRSS) equations [3] were adapted to 37°C [29] and used to describe non linear membrane dynamics. The internodal myelin was considered as a perfect insulator. Axonal activation was defined by the induction and propagation of action potentials (AP) along at most 10 nodes of Ranvier. The model included 15808 fibers with 41 nodes of Ranvier.

### Stimulation parameters

Monophasic versus biphasic stimulation applied with monopolar and bipolar electrodes (Fig. 1) were compared. Stimulus parameters were chosen in accordance with studies that record axono-cortical evoked potentials (ACEP; *PW* = 500-1000 μs, *I*= 0.5 *mA* to 10 *mA*). The stimulus duration was 500 μs for monophasic stimulation. In case of monopolar stimulation, cathodic (depolarizing) stimuli were only applied to the electrode placed on the axonal tract whereas anodic (hyperpolarizing) stimuli were applied to the distant contact (reference). Biphasic stimulation was considered without inter-stimulation delay. Each phase lasts 500 μs for a total biphasic stimulation of *PW* = 1000 μs. All electrode and stimulation parameters are listed in Table. 1. Electrode orientations were placed orthogonal (transverse) or parallel to the axonal tact (Fig. 1). The values for stimulus amplitude *I* were 0.5, 1, 1.5, 2, 2.5, 3, 5 and 10 *mA*.

**Table 1.** Electrode configuration, stimulus shape and axonal tract conductivity used in this study.

| Electrode parameter | Stimulus shape | Fasciculus conductivity |
| --- | --- | --- |
| monopolar | monophasic | isotropic |
| monopolar | biphasic | isotropic |
| monopolar | monophasic | anisotropic |
| monopolar | biphasic | anisotropic |
| bipolar orthogonal | monophasic | isotropic |
| bipolar orthogonal | biphasic | isotropic |
| bipolar orthogonal | monophasic | anisotropic |
| bipolar orthogonal | biphasic | anisotropic |
| bipolar parallel | monophasic | isotropic |
| bipolar parallel | biphasic | isotropic |
| bipolar parallel | monophasic | anisotropic |
| bipolar parallel | biphasic | anisotropic |

### Evaluation of axon activation

Maximal activation depth and area were evaluated as functions of electrode parameters (monopolar, bipolar orthogonal, or bipolar parallel), stimulus shape (monophasic or biphasic), electrical conductivities (isotropic or anisotropic), and stimulus amplitude. The maximal activation depth is defined as the distance between the electrode and the most deeply activated axon. The stimulation area quantifies the white matter surface activated by direct stimulation (cathode) or indirect stimulation (virtual cathodes).

## Results

### Action potential propagation: effect of pulse shape

We first studied the effect of pulse shape on action potential (AP) propagation (Fig. 2 A-C). Several phenomena result from bipolar orthogonal monophasic stimulation: activation by the cathode (Fig. 2 C-D), an activation under the anode due to virtual cathodes also called anode-make stimulation [25] (Fig. 2 C,E), and AP blocking under the cathode following initiation of an AP, but absence of propagation between the nodes of Ranvier (Fig. 2 C,F). Such a blocking phenomenon appears for axons very close to the cathode [23]. Therefore, only axons located in a shell around the electrode are stimulated (Fig. 3). AP blocking was mainly observed for monophasic stimulation regardless the stimulus amplitude and to a lesser extent for biphasic stimulation with amplitude greater than 5 *mA*.

**Figure 2.**
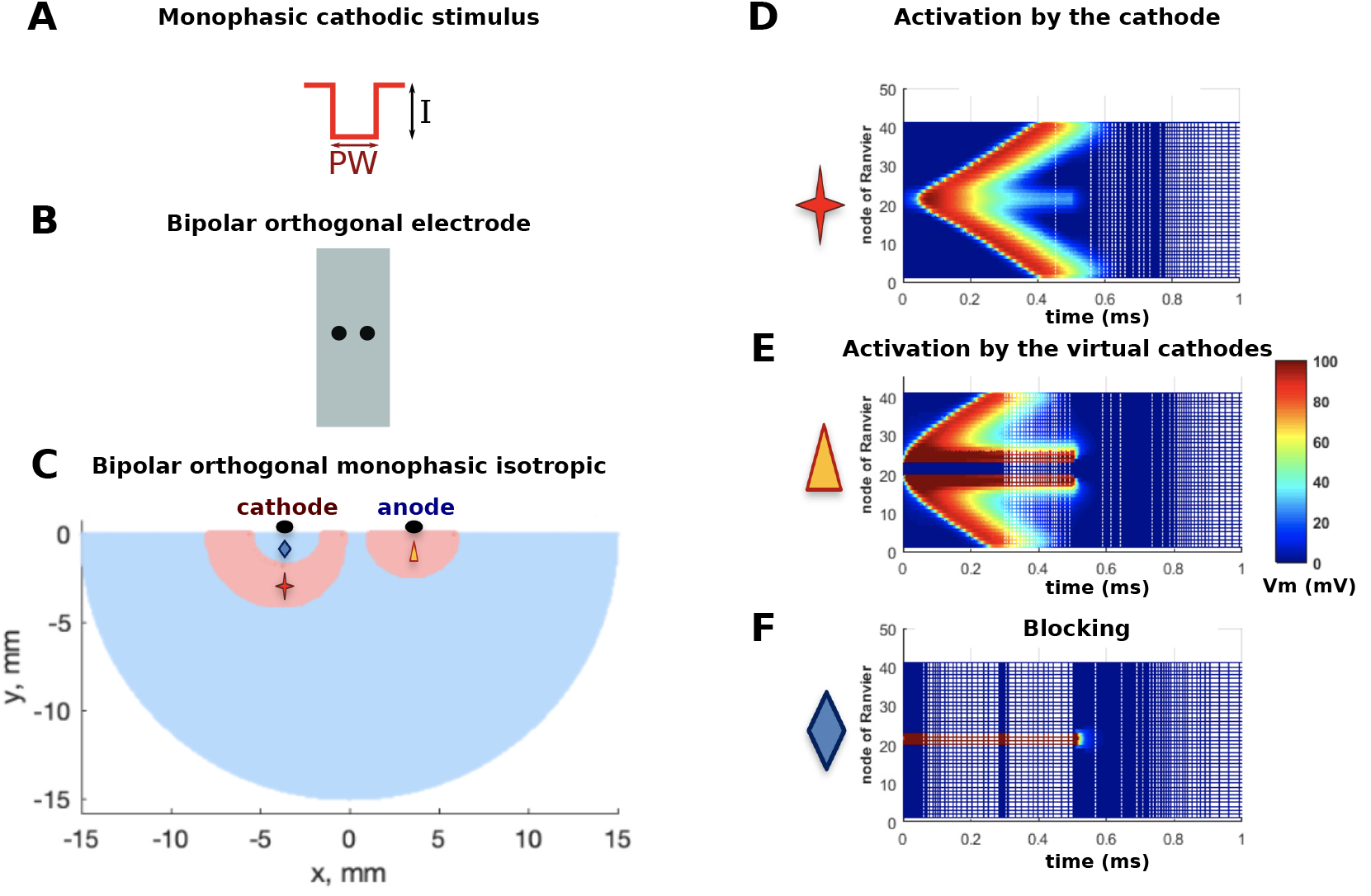
AP propagation or blocking along Nodes of Ranvier. (A-C): Example stimulation: bipolar orthogonal electrode with monophasic stimulus (1 *mA*). (D): Activation and AP propagation by direct cathodic stimulus, (E): Activation and AP propagation by virtual cathode, (F): AP is blocked. *V*_*m*_ is the membrane voltage of the axon.

**Figure 3.**
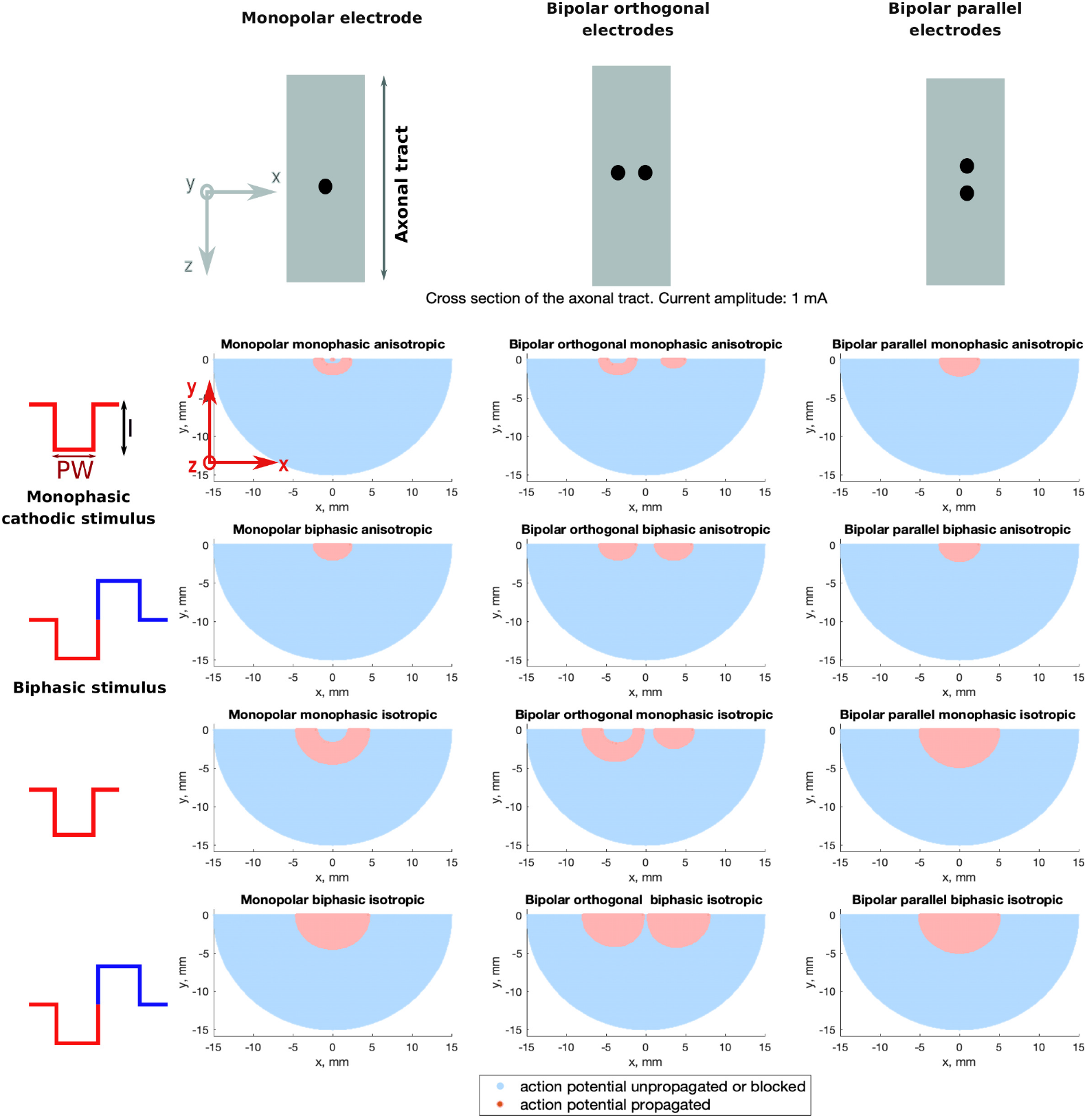
Map of stimulus effects. Cross-section of axonal tacts, representing axons activated, inactivated, or blocked by electrical stimulation. Stimulus amplitude: 1 *mA*.

### Activation map

The stimulated area is shown on cross-sections of the axonal tracts (Fig. 3, stimulus amplitude: 1 *mA*) for all the configurations tested. At this amplitude, for anisotropic tracts, maximal activation depth is similar giving 1.8 *mm* for monopolar and bipolar orthogonal configurations and 2 *mm* for bipolar parallel configuration. Considering isotropic tracts, maximum activation depth was 4.7 *mm* for the bipolar parallel configuration, 4.3 *mm* for the monopolar configuration and 3.9 *mm* for the bipolar orthogonal configuration. The activation map at 10 *mA* is shown in appendix Fig. 7. Note that at this amplitude, a blocking effect appears using biphasic stimulus. Thus, applying biphasic stimulus, maximal activation depth occurred under the both contacts of the electrode, whereas maximal activation depth occurred under the cathode in case of monophasic stimulation. Activation under the anode (due to virtual cathodes) was observed using monophasic stimulus.

### Influence of the stimulus amplitude

#### Anisotropic model

Considering an anisotropic axonal tract model as a function of the stimulus amplitude, Fig. 4A shows the maximal activation. As expected, the maximal activation depth increased with current amplitude. No difference of maximal activation depth was found between monophasic and biphasic stimulus. Activation depth difference between monopolar and bipolar orthogonal or bipolar parallel configuration was 0.1 *mm* below 10 *mA*. Since the points of the axon grid are separated by 0.1 *mm*, this difference was not considered significant. Maximum activation depth was reached using monopolar configuration at 10 *mA* (4.6 *mm*). Activation depth difference between monopolar and bipolar orthogonal or bipolar parallel was 0.3 *mm*. Note that the monopolar and bipolar parallel curves intersect at 5 *mA*.

**Figure 4.**
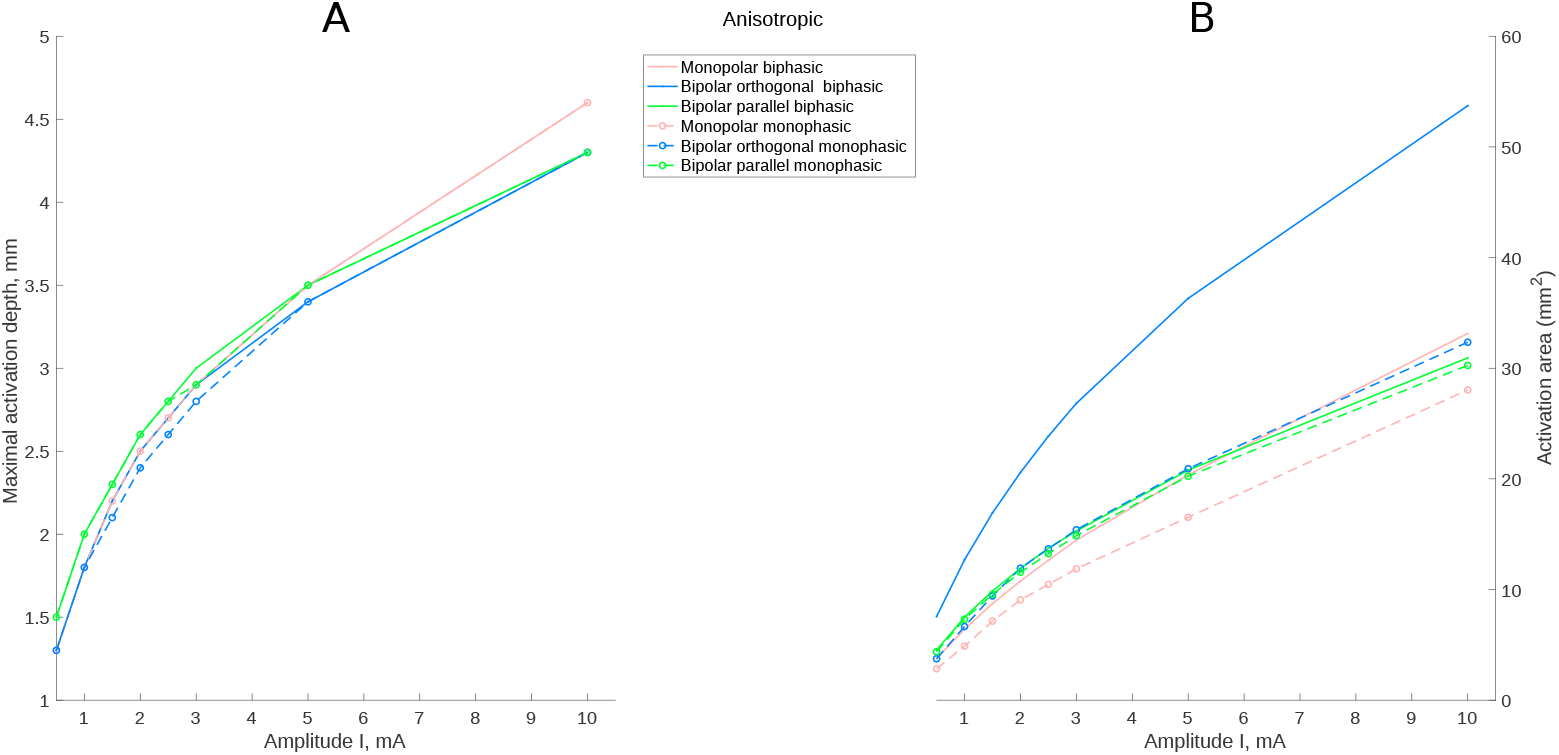
Anisotropic model. (A): Maximal activation depth and (B): activation area in function of the stimulus amplitude

The Fig. 4B, shows the variation of the activation area in function of stimulus amplitude. The largest activation area was obtained with bipolar biphasic stimulation (7.55 *mm*^2^ at 0.5 *mA* to 53.7 *mm*^2^ at 10 *mA*). Compared to monophasic stimulation, the biphasic stimulation recruited 79 %, 24 % and 3% more fibers respectively for bipolar orthogonal, monopolar, and bipolar parallel configurations respectively. The monopolar monophasic configuration recruited the smallest area (2.84 *mm*^2^ at 0.5 *mA* to 28.0 *mm*^2^ at 10 *mA*).

#### Isotropic model

The same analysis was performed considering the axonal tract isotropic. No difference of maximal activation depth were found between monophasic and biphasic stimulus (Fig. 5A). However, the maximal activation depth difference was significant between monopolar, bipolar orthogonal, and bipolar parallel configurations. The maximal depth difference between monopolar and bipolar orthogonal increased linearly with increasing current amplitude I (*y* = 0.1955 × *I* + 0.1144, *R*^2^ = 0.99). The maximal depth difference between bipolar parallel and monopolar was stable (0.5 *mm*) from 1 *mA* to 5 *mA*. At 10 *mA* the monopolar and bipolar parallel curves intersect. Maximum activation depth was reached using monopolar and bipolar parallel configurations at 10 *mA* (10.7 *mm*).

**Figure 5.**
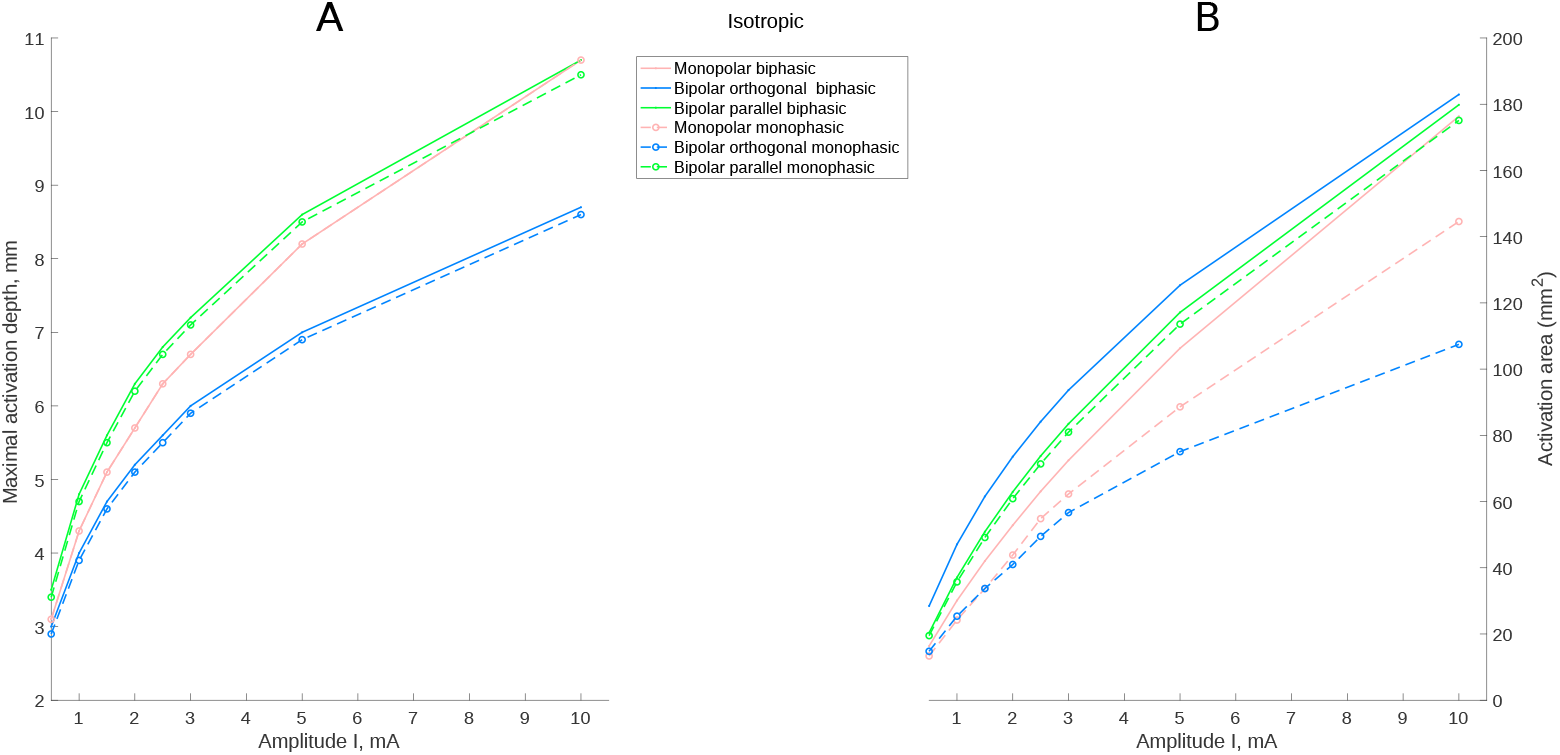
Isotropic model. (A): Maximal activation depth and (B): activation area in function of the stimulus amplitude

The variation of activation area as a function of stimulus amplitude is shown in Fig. 5B. The largest activation area was obtained with bipolar biphasic stimulation (28.5 *mm*^2^ at 0.5 *mA* to 182.9 *mm*^2^ at 10 *mA*). Compared to monophasic stimulation, the biphasic stimulation recruited 76 %, 21 % and 3% more fibers respectively for bipolar orthogonal, monopolar, and bipolar parallel configurations. The monopolar monophasic configuration activated the smallest area from 1 *mA* to 1.5 *mA* whereas from 2 *mA* to 10 *mA*, the bipolar monophasic configuration performed more poorly.

#### Influence of conductivity on the area stimulated

Comparison between isotropic and anisotropic case concerning maximal activation depth and activation area are shown in Fig. 6 considering a biphasic pulse. Using isotropic model overestimated the maximal activation depth and the activation area compared to the anisotropic model. In averaged, activation area was 257 %, 391 % and 427 % more important respectively for bipolar orthogonal, monopolar and bipolar parallel configuration. The difference increased with current amplitude for the monopolar and bipolar parallel configuration but decreased for the bipolar orthogonal configuration. The maximal activation depth was 112 %, 134 % and 142 % more important respectively for bipolar orthogonal, monopolar and bipolar parallel configuration.

**Figure 6.**
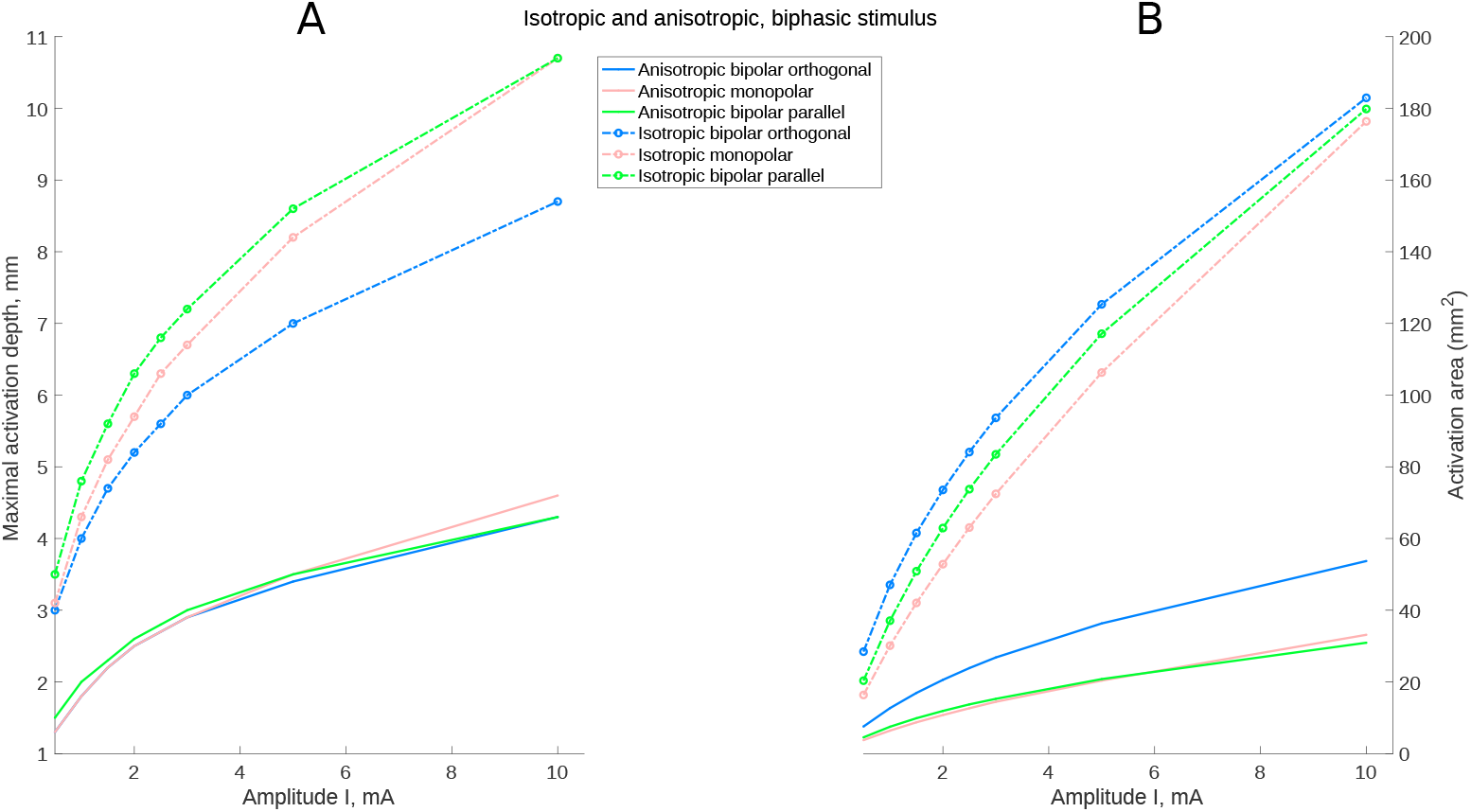
Comparison between isotropic and anisotropic model, using biphasic stimulus. (A): Maximal activation depth and (B): activation area in function of the stimulus amplitude.

## Discussion

In this study, we report the investigation of parameters used in surgical electrical stimulation of white matter tracts using computational models. The results show that different electrode parameters (bipolar parallel or orthogonal electrode, monopolar electrode, biphasic or monophasic stimulus) drastically modify the area and depth of tract activation.

Biphasic bipolar orthogonal stimulation is widely used in surgery [31] and activates a larger total area than monopolar or bipolar parallel stimulation, however, axons lying between bipolar electrodes remain inactive. These results are in agreement with a previous study modelling white matter as a continuous bidomain medium [17]. Specifically, activation areas symmetrically surround the two poles of the bipolar biphasic orthogonal probe, rather than being located in between the two poles. This is in line with the fact that it is the second spatial derivative of the potential that drives the membrane depolarization (see the notion of “activating function” [15, 24]). Hence, from a practical point of view, the effect of a biphasic bipolar orthogonal stimulation can be approximated as a double biphasic monopolar stimulation.

Consistent with the recent report of Gomez et al. for isotropic tracts, monopolar, monophasic stimulation caused a broader and deeper activation than monophasic stimulation with bipolar electrodes orthogonal to axon tracts [8]). However, our results show that monophasic stimulation with bipolar electrodes parallel to isotropic tract is far more effective than either monopolar or bipolar orthogonal electrodes. Further, the use of monophasic stimulation can induce potentially troublesome phenomena such as virtual cathodic activation (under the anode) and blocking of AP under the cathode. Notably, a non-symmetrical stimulation occurs below the contacts of the bipolar orthogonal electrode such that the stimulation zone is less well controlled.

The spatial features of axon recruitment by the different configurations were profoundly affected by anisotropy, both qualitatively and quantitatively. Assuming an anisotropic axonal tract, we found that there was no difference in activation depth using monopolar or bipolar configurations at low stimulus amplitude whereas the difference is clearly visible in an isotropic tract. The maximal depth and total activation area were far greater in isotropic compared to anisotropic models. The common result in both anisotropic and isotropic models was that bipolar orthogonal biphasic stimulation activates a larger total area than monopolar or bipolar parallel stimulation. Our findings indicate that inhomogeneities of conductivity have a drastic effect on the area recruited by the stimulation. It is thus important to precisely map the conductivities experimentally in order to enable the design of precise models. By extension, possible non-ohmic properties of the extracellular space [7] could also influence the area stimulated. In future studies, realistic geometrical head models including realistic conductivities, fiber densities, and inhomogeneities of axon diameter should be considered to construct patient- and tract-specific predictions.

## Acknowledgment

Funding: This work was suported by the CRC chirurgie 2016 from AP-HP, INSERM through Interface Contracts for Hospitals, CNRS, and the European Community (Human Brain Project H2020-785907).

## Appendix

**Figure 7.**
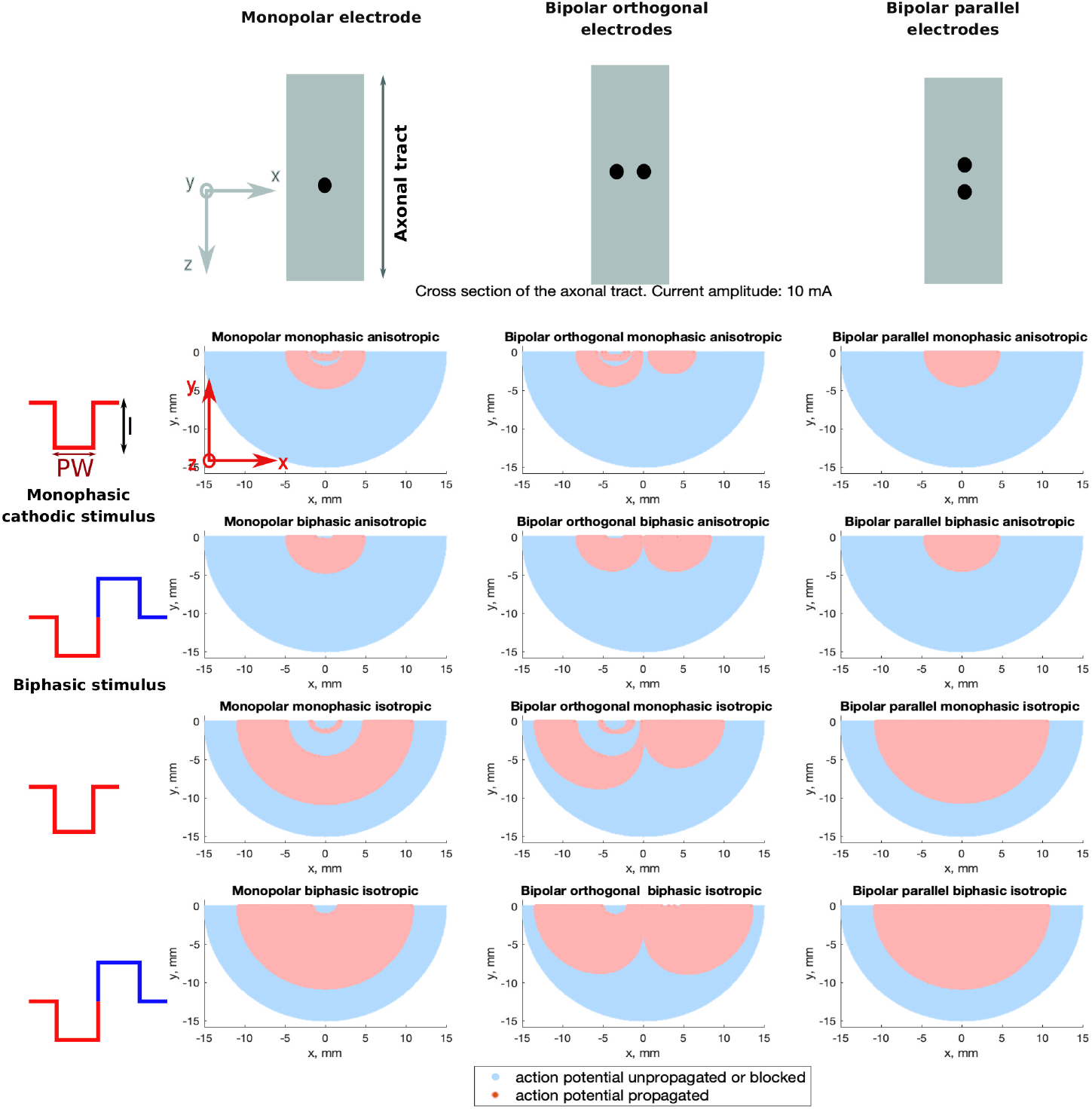
Map of stimulus effects. Cross-section of axonal tacts, representing axons activated, inactivated, or blocked by electrical stimulation. Stimulus amplitude: 10 *mA*.

